# Multiscale communication in cortico-cortical networks

**DOI:** 10.1101/2020.10.02.323030

**Authors:** Vincent Bazinet, Reinder Vos de Wael, Patric Hagmann, Boris C. Bernhardt, Bratislav Misic

**Affiliations:** McConnell Brain Imaging Centre, Montréal Neurological Institute, McGill University, Montréal, Canada; Department of Radiology, Lausanne University Hospital (CHUV-UNIL), Lausanne, Switzerland

## Abstract

Signaling in brain networks unfolds over multiple topological scales. Areas may exchange information over local circuits, encompassing direct neighbours and areas with similar functions, or over global circuits, encompassing distant neighbours with dissimilar functions. Here we study how the organization of cortico-cortical networks mediate localized and global communication by parametrically tuning the range at which signals are transmitted on the white matter connectome. By investigating the propensity for brain areas to communicate with their neighbors across multiple scales, we naturally reveal their functional diversity. We find that unimodal regions show preference for local communication and multimodal regions show preferences for global communication. We show that these preferences manifest as region- and scale-specific structure-function coupling. Namely, the functional connectivity of unimodal regions emerges from monosynaptic communication in small-scale circuits, while the functional connectivity of transmodal regions emerges from polysynaptic communication in large-scale circuits. Altogether, the present findings reveal how functional hierarchies emerge from hidden but highly structured multiscale connection patterns.

## INTRODUCTION

The brain is a network of anatomically connected neuronal populations [17]. This complex web of connections functions as a communication network, promoting signaling between brain regions [3, 33]. A tendency for neuronal populations with similar functions to connect with each other gives rise to a nested hierarchy of increasingly polyfunctional neural circuits, spanning multiple topological scales [41, 47, 95].

Studies of network communication typically conceptualize signalling events as a global process, eschewing the possibility that communication takes places over multiple topological scales. Namely, areas may preferentially exchange information over small compact circuits encompassing direct neighbours and areas with similar functions, or over more extensive circuits encompassing more distant neighbours with dissimilar functions. An intuitive example is the worldwide air transportation network. The purpose of regional or domestic flights permitting transit between a country’s regions is different from the purpose of international flights permitting transit between international hubs. The importance of an airport in this network will correspondingly depend on the type of flight considered. For example, Denver’s and Philadelphia’s airports are important for domestic flights within the United States, while airports in New York, Los Angeles or Chicago are important for international flights [36]. In other words, the topological role of a node in the network depends on the scale at which it is evaluated.

By the same token, individual brain areas may exhibit characteristic interactions and communication patterns at multiple topological scales. The modular structure of the brain [40], in concert with a prominent connective core of high degree areas [80, 81], creates conditions in which information can be either segregated into local clusters of highly interconnected brain regions, or globally integrated [93, 94]. For instance, an area may facilitate the integration of information among its local neighbours, but lack the capacity to globally broadcast signals across the whole brain. In other words, the functional diversity of a region – who it can communicate or interact with – depends on scale.

Here we study how communication between brain regions unfolds over multiple scales. For a given region, we systematically define local neighborhoods of increasing size. We then track how the centrality of individual brain regions varies as the sizes of the probed neighborhoods increase. We show that variations in centrality are shaped by functional diversity. We further find a localized-distributed gradient of communication preferences, such that unimodal regions are preferentially central locally and transmodal regions are preferentially central globally. Finally, we demonstrate that structure-function coupling is scale-specific, such that the functional connectivity profiles of unimodal regions are better captured by communication within small-scale structural neighbourhoods, while the functional connectivity profiles of transmodal regions are better captured by communication within large-scale structural neighbourhoods.

## RESULTS

The results are organized as follows. We delineate multiscale neighborhoods by parametrically tuning the range at which signals are transmitted on the white matter connectome. We subsequently investigate the propensity for brain areas to communicate with their neighbours across multiple scales using a weighted measure of regional closeness centrality. Finally, we consider how the similarity in two areas’ embedding predicts their functional connectivity. Data sources include (see *Materials and Methods* for detailed procedures):

- *Structural connectivity*. Structural connectomes were generated for *N* = 67 healthy participants (source: Lausanne University Hospital). Individual weighted network were reconstructed using diffusion spectrum imaging and deterministic streamline tractography.
- *Functional connectivity*. Functional connectivity was estimated in the same individuals (*N* = 67) using resting-state functional MRI (rs-fMRI).

Analyses were performed using a network parcellation of 1000 cortical nodes [18]. They were subsequently repeated using coarser resolutions (114, 219 and 448 nodes) and an independently collected dataset (HCP; *N* = 201) (see *Materials and Methods* for more information on the *Validation* dataset).

### Multiscale regional centrality

We first characterize local neighborhoods, in each structural connectome, using unbiased random walks. Specifically, we use the transition probabilities of a random walker seeded in an individual brain region to delineate its local neighborhood (see *Methods* for more details). Transition probabilities were measured for 100 time scales *t*, logarithmically spaced between 10^*–*0.5^ and 10^1.5^. Fig. 1a shows the effect of varying the scale of a random walk initiated at nodes located in the posterior cingulate (red), superior parietal (blue), transverse temporal (green) and insular (purple) cortices, with *t* = 2, *t* = 5 and *t* = 10. As *t* is increased, the random walks are longer and the size of the probed neighborhood becomes larger, allowing us to consider communication over more expansive portions of the network.

**Figure 1.**
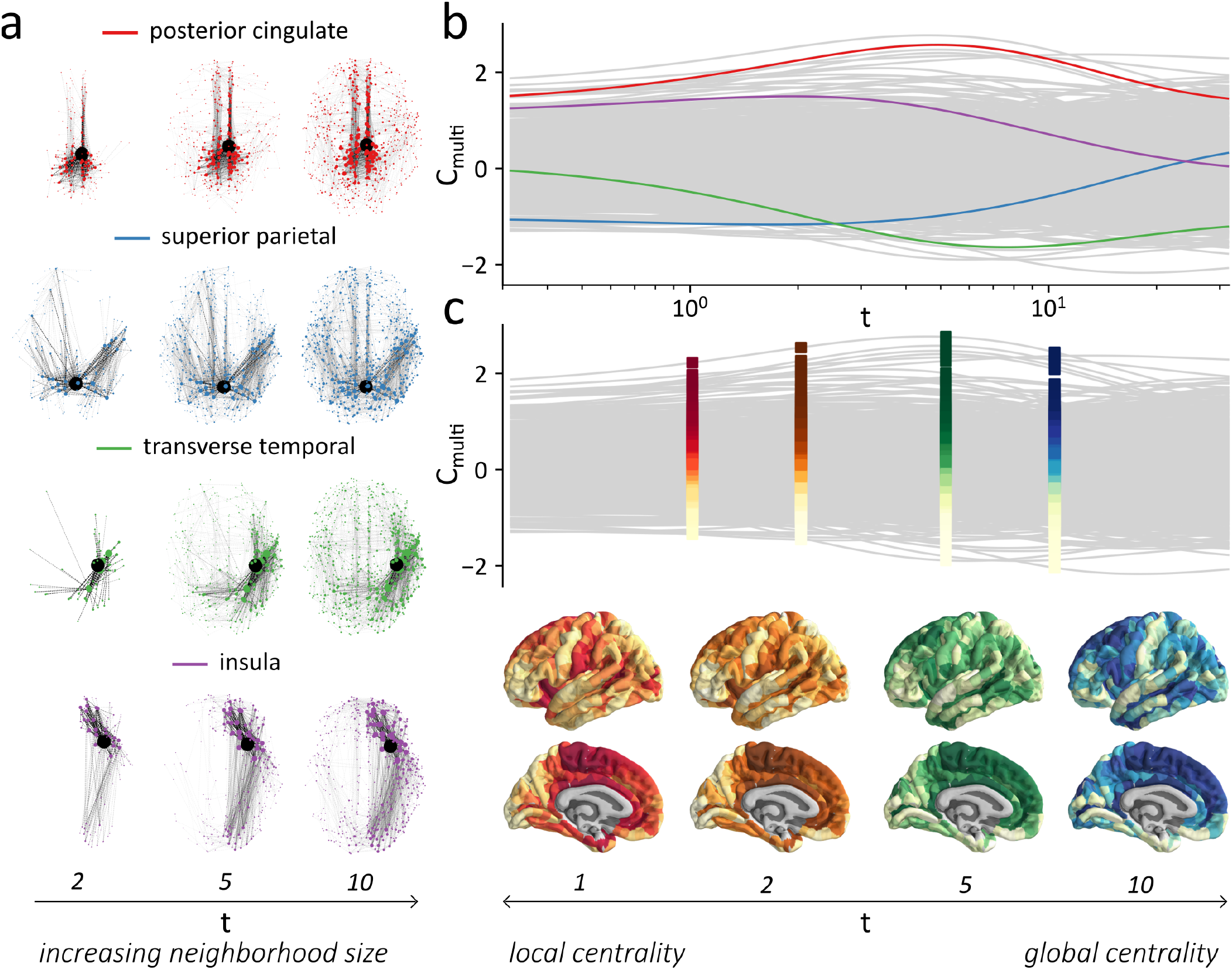
Multiscale regional centrality. **(a)** An unbiased random-walk process can be used to delineate local neighborhoods around individual brain regions of the structural connectome. As *t*, the number of iterations, is increased, the topological size of the characterized neighborhoods gets larger. Random walk process can be initiated, for example, in the posterior cingulate gyrus, (red), in the superior parietal gyrus (blue), in the transverse temporal gyrus (green) or in the insula (purple), and can delineate neighborhoods of different sizes (*t* = 2; *t* = 5, *t* = 10). **(b)** The weights of a node’s transition probability vector are used to compute a multiscale measure of closeness centrality (*C*_multi_; see *Methods* for more details). Grey lines represent the centrality of individual brain regions, averaged across subjects, as *t* increases. The centrality scores are standardized for each time scale *t*. Highlighted in red, blue, green and purple are the centrality trajectories of the four individual brain regions shown in (a). A node’s relative centrality varies largely depending on the scale of the neighborhoods. **(c)** The relative importance of a brain region in local communication processes can be evaluated by ranking *C*_multi_ scores within small neighborhoods (*t* = 1; left-most). Its relative importance in global processes (global centrality) can be evaluated by ranking *C*_multi_ scores within large neighborhoods (*t* = 10, right-most). The importance of a brain region in communication processes unfolding at intermediate scales can be similarly evaluated (e.g. *t* = 2 or *t* = 5). Darker colors indicate brain regions with large *C*_multi_ ranks while lighter colors indicate brain regions with low *C*_multi_ ranks.

To investigate how the role of different brain regions varies across scales, we measure a region’s closeness to other nodes in its local neighborhood. Closeness centrality is typically computed as the average of local scores measuring the inverse of the shortest path between a region of interest and each individual region in the network. Here we weight the local scores, for individual nodes, according to their proximity to the region of interest using the transition probability vector as a weight function (see *Methods*). This weight function reflects the intuition that the greater the number of electrochemical synapses a signal has to traverse, the greater the conductance time and potential attenuation of that signal [27]. When studying local interactions between proximal neuronal populations, distant neighbors become less relevant because signals cannot reach them as readily [79]. By measuring how easily a brain region can communicate with neighbors characterized across different topological scales, we obtain a multiscale measure of closeness centrality (*C*_multi_).

For each time scale, we computed the *C*_multi_ of every brain region. Fig. 1b shows regional values of *C*_multi_ as *t* increases, averaged across subjects. To allow for comparisons between scales, *C*_multi_ scores are standardized relative to the distribution of scores obtained at individual scales. The centrality trajectories of four sample brain regions are highlighted, and others are shown in grey. The relative centrality of individual brain regions varies considerably with increasing scale, such that some areas are more central, and some are less central, depending on the size of their neighbourhood. Fig. 1c illustrates the centrality of every brain region, averaged across subjects, for four different topological scales. Locally, we observe clusters of highly central brain regions distributed across the whole brain; the clusters gradually evolve into larger-scale systems at more global scales.

### Multiscale functional diversity

In homogeneous networks where nodes have similar topological characteristics, the local centrality of a node is expected to be similar to its global centrality. However, in heterogeneous networks, local attributes do not necessarily mirror global attributes [25]. For instance, one node may have strong connectivity with a small number of nodes, while another node may have moderate but diverse connectivity with a larger number of nodes. The former is more central in a local sense, while the latter is more central in a global sense. The differential contribution of the two nodes to global communication, arising from their respective functional diversity, is reflected by their different closeness trajectories. See Fig. S1 for an illustration of this concept in an artificially generated network.

To quantify variations in a brain region’s centrality across scales, we compute the local *slopes* in the closeness trajectories of individual nodes of the structural connectomes as *t* increases. Fig. 2a shows the *C*_multi_ trajectories of four nodes located in the posterior cingulate, superior parietal, transverse temporal and insular cortices (from Fig. 1a,b), colored according to their slope. By considering the slope of a brain region at a single scale, we can measure its functional diversity, with highly diverse brain regions having a positive slope (red) and less diverse brain regions having a negative slope (blue). Fig. 2b shows how the topographic distribution of these slopes on the brain varies across scales. Importantly, these slopes capture the “functional diversity” of a brain region rather than its “connection diversity”, because it considers the diversity of a node’s relationships across a neighborhood of arbitrary size, as opposed to the diversity of its direct connections.

**Figure 2.**
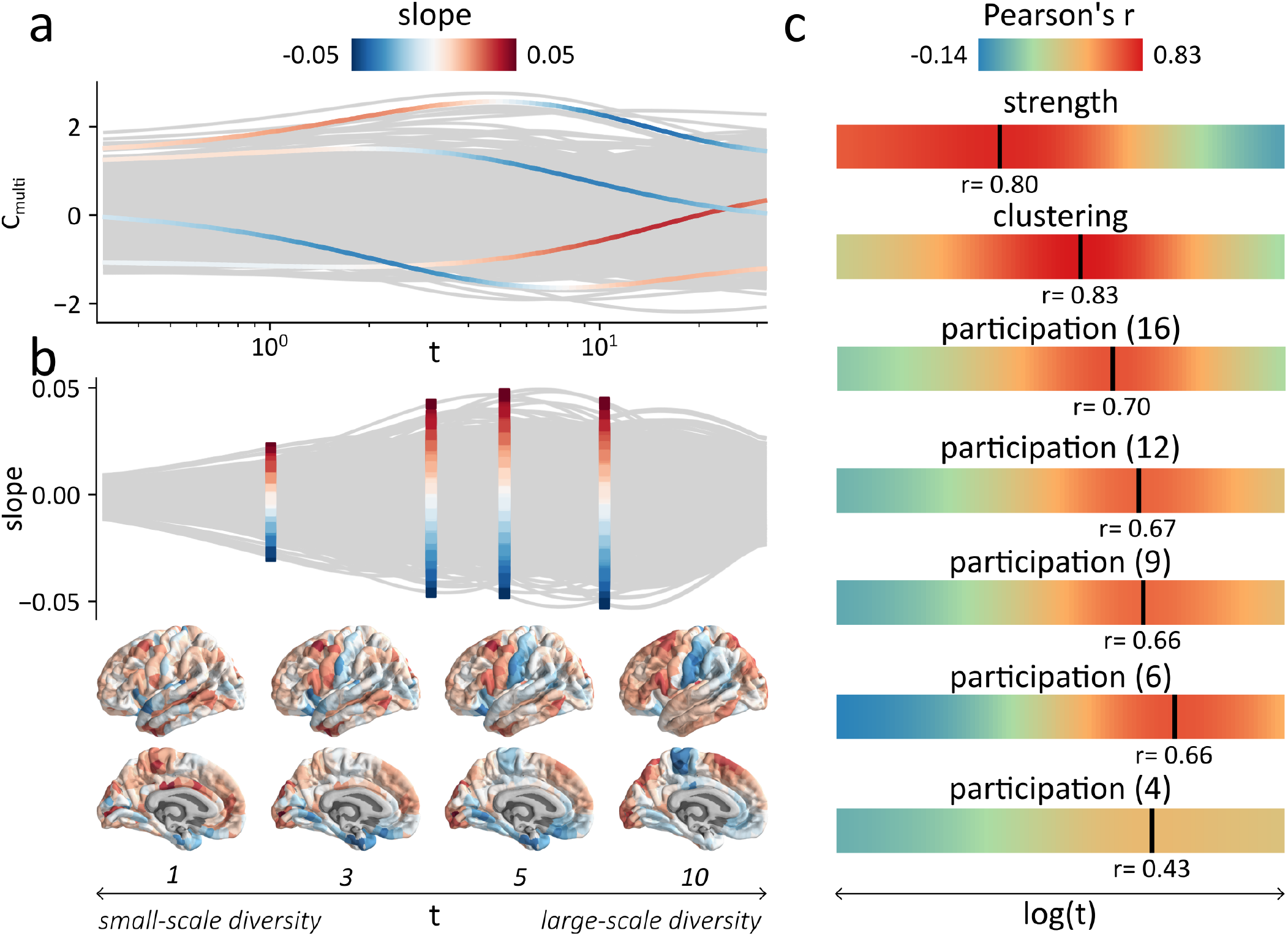
Multiscale functional diversity. **(a)** The functional diversity of a brain region is quantified as the amplitude of the local variations in a region’s closeness centrality (*slope*) as *t* varies. **(b)** Slopes can show different topographic distribution, for select values of *t*. Highly diverse brain regions have a positive slope (red) while less diverse brain regions have a negative slope (blue). **(c)** Slope scores are correlated, as *t* increases, with other measures of connection diversity (node degree, clustering coefficient, participation coefficients for partitions of 16, 12, 9, 6 and 4 communities). Local measures of diversity such as degree and clustering coefficient (negative) are correlated with the centrality slopes measured at intermediate scales, with peaks at *t* = 1.69 (r=0.80) for node degree and *t* = 3.90 (r=0.83) for clustering coefficient (negative). Participation coefficients, viewed as meso-scale measures of functional diversity, are also correlated with a node’s local variation in centrality. The scale at which the correlation peaks highlights the size of the communities used to compute the participation coefficient, with larger values of *t* measuring the role of brain regions inside larger communities.

To demonstrate how transitions in local closeness can highlight the functional diversity of a brain region at multiple scales, we computed seven measures capturing a node’s connection diversity at different topological scales. These measures are nodal strength, clustering coefficient and participation coefficients computed with respect to modular partitions of the networks into 16, 12, 9, 6 and 4 communities. For each of the seven measures, we averaged the scores obtained across subjects and correlated them with the *C*_multi_ slopes. Fig. 2c shows the correlations between these measures and the local slopes evaluated across time scales. Measures are ordered from top to bottom according to the scale at which they are maximally correlated with the closeness slope. Local measures, such as strength and clustering coefficient, tend to be maximally correlated at lower scales (*t* = 1.69 and *t* = 3.90, respectively). Participation coefficients – indexing the diversity of inter-modular links for a given partition – tend to be correlated with local variations in closeness at greater values of *t*. Furthermore, as the partition resolution is gradually decreased, from 16 to 4 communities, optimal correlations are obtained at larger values of *t*. Altogether, these results demonstrate that variations in a node’s centrality are mediated by its functional diversity. Interestingly, the present method serves to highlight functional diversity without a predefined partition, making it a complementary measure to more traditional diversity statistics (such as the participation coefficient), without the need to explicitly define or assume a hard partition.

### Optimal communication scales

What is the optimal scale at which individual brain regions communicate? Fig. 3a shows the scale (*t*) at which the centrality of individual brain regions in the structural connectomes peaks (*t*_opti_). The *t*_opti_ values were averaged across subjects. Cooler colors indicate regions that preferentially communicate locally while warmer colors indicate regions that preferentially communicate globally. In general, we observe preference for local communication in primary sensory regions (pericalcarine cortex, transverse temporal cortex, post-central gyrus) and in the limbic cortex; conversely, we observe preference for global communication in association cortex, including dorsolateral prefrontal cortex and superior parietal cortex.

**Figure 3.**
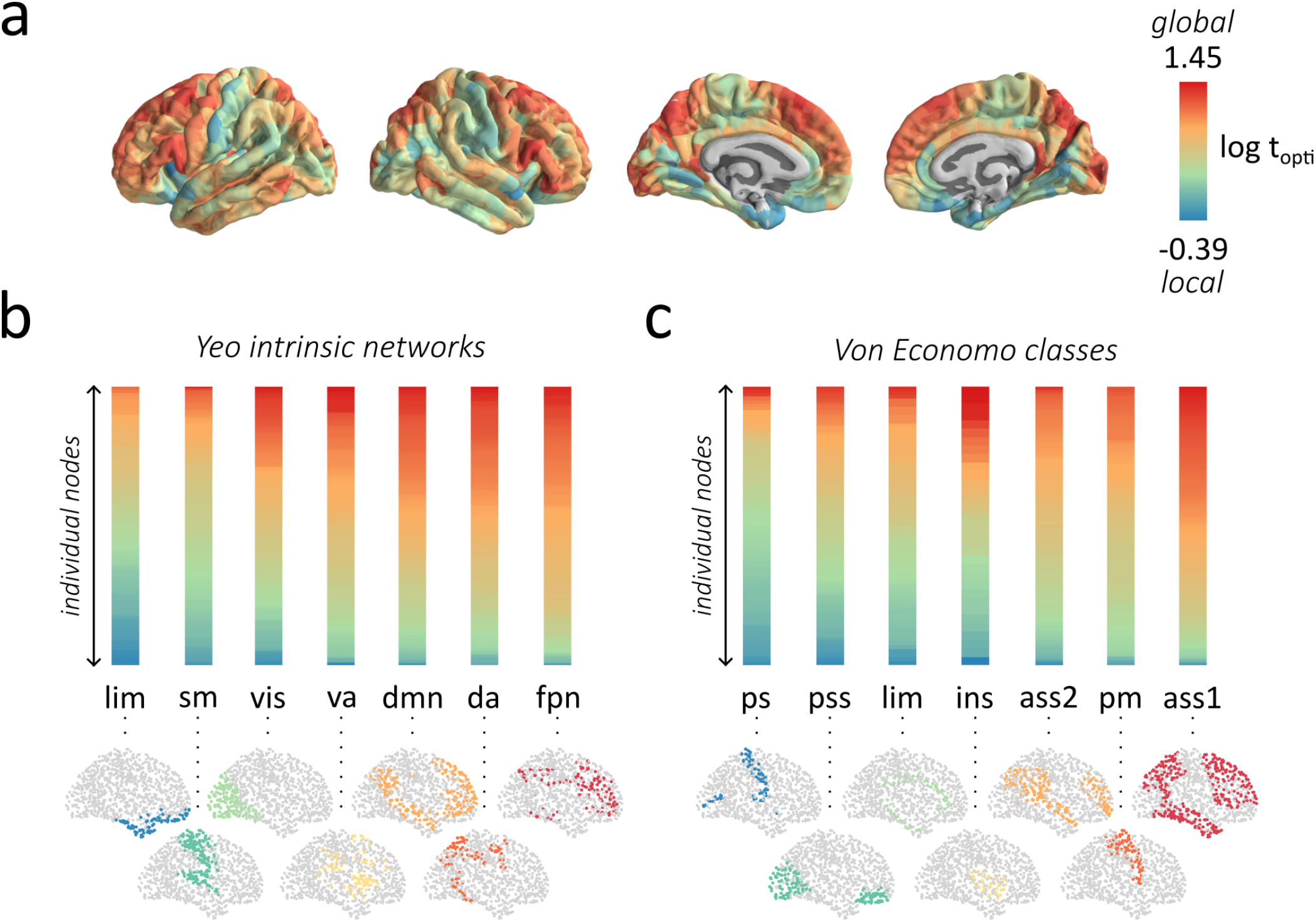
Optimal communication scales. **(a)** Representation on the surface of the brain of the optimal scale (*t*_*opti*_) at which the multiscale closeness centrality of individual brain regions of the structural connectome peaks. The *t*_*opti*_ scores are averaged across participants. Sensory region are optimally central for low values of *t* and multimodal regions are optimally central for large values of *t*. **(b)** Heatmaps of the distribution of optimal scale values for individual brain regions, averaged across subjects, for seven intrisic functional networks [91]. The nodes in each heatmap are vertically ordered and colored based on their *t*_*opti*_ scores. **(c)** Heatmaps of the distribution of optimal scale values for individual brain regions, averaged across subjects, for seven cytoarchitectonic classes defined from the von Economo atlas [84, 86–88]. Yeo intrinsic networks: lim = limbic network, sm = somatomotor network, vis = visual network, va = ventral attention network, dmn = default mode network, da = dorsal attention network, fpn = frontoparietal network. Von Economo classes: ps = primary sensory cortex, pss = primary/secondary sensory cortex, limbic = limbic cortex, insular = insular cortex, ass2 = association cortex 2, pm = primary motor cortex, ass1 = association cortex 1.

Fig. 3b shows the distribution of optimal values of *t* for seven intrinsic functional networks [91]. These distributions are represented as heatmaps such that the nodes in the intrinsic functional networks are vertically ordered and colored based on their *t*_opti_ values. The mean of the distributions for the seven networks are significantly different from one another following Bonferroni correction (p<0.001), except for the mean *t*_opti_ scores of the dorsal attention and default-mode networks (p=0.45) and for the dorsal attention and fronto-parietal networks (p=0.04). These heatmaps highlight a differentiation between limbic and unimodal (somatomotor and visual) networks versus multimodal networks (default-mode, dorsal-attention and fronto-parietal networks). Fig. 3c shows, in the same way, the distribution of *t*_opti_ values for seven cytoarchitectonic classes defined by the von Economo atlas [84, 86–88].The means of the distributions for the primary sensory (ps) and association (ass1) classes are significantly different from the mean of the other six cytoarchitectonic classes following Bonferroni correction (p<0.001). These heatmaps again highlight a differentiation between sensory and association areas.

We also compared the averaged *C*_multi_ of the seven intrinsic functional networks and cytoarchitectonic classes, as *t* increases, to the average *C*_multi_ of these intrinsic networks and classes in randomized networks with preserved degree sequences (Fig. S2). We find that the variations in *C*_multi_ are larger in the empirical networks than those in the randomized networks. In other words, variations in the centrality of a node are not trivially expected from their degree.

### Multiscale structure-function coupling

We next investigate how multiscale connection patterns influence structure-function coupling. Functional connectivity between pairs of brain regions is typically computed as a correlation between the time series of their respective fMRI BOLD signals. Coherent fluctuations in neural activity are thought to arise from interactions on the underlying structural connectome [9, 32, 75]. A variety of pair-wise measures have been proposed to predict functional connectivity from the structural connectivity between brain regions, including structural connectivity strength, path length, search information, path transitivity [32] and communicability [9]. Most of the proposed measures of structure-function coupling assume a single-scale, global relationship between the two (but see [1, 9] for their use of multiscale measures).

Here we assess structure-function coupling across multiple scales. By measuring the similarity between the neighborhoods of two brain regions defined at different values of *t*, we ask how similar the dynamical processes unfolding around them are [70]. We hypothesize that nodes with overlapping neighborhoods (i.e. large neighborhood similarity) will display greater functional coupling than nodes that are part of different neighborhoods [56]. We hence quantified, for each structural connectome, the neighborhood similarity of pairs of brain regions by computing the pairwise cosine similarity between their transition probability vectors, for every value of *t* (Fig. 4a).

**Figure 4.**
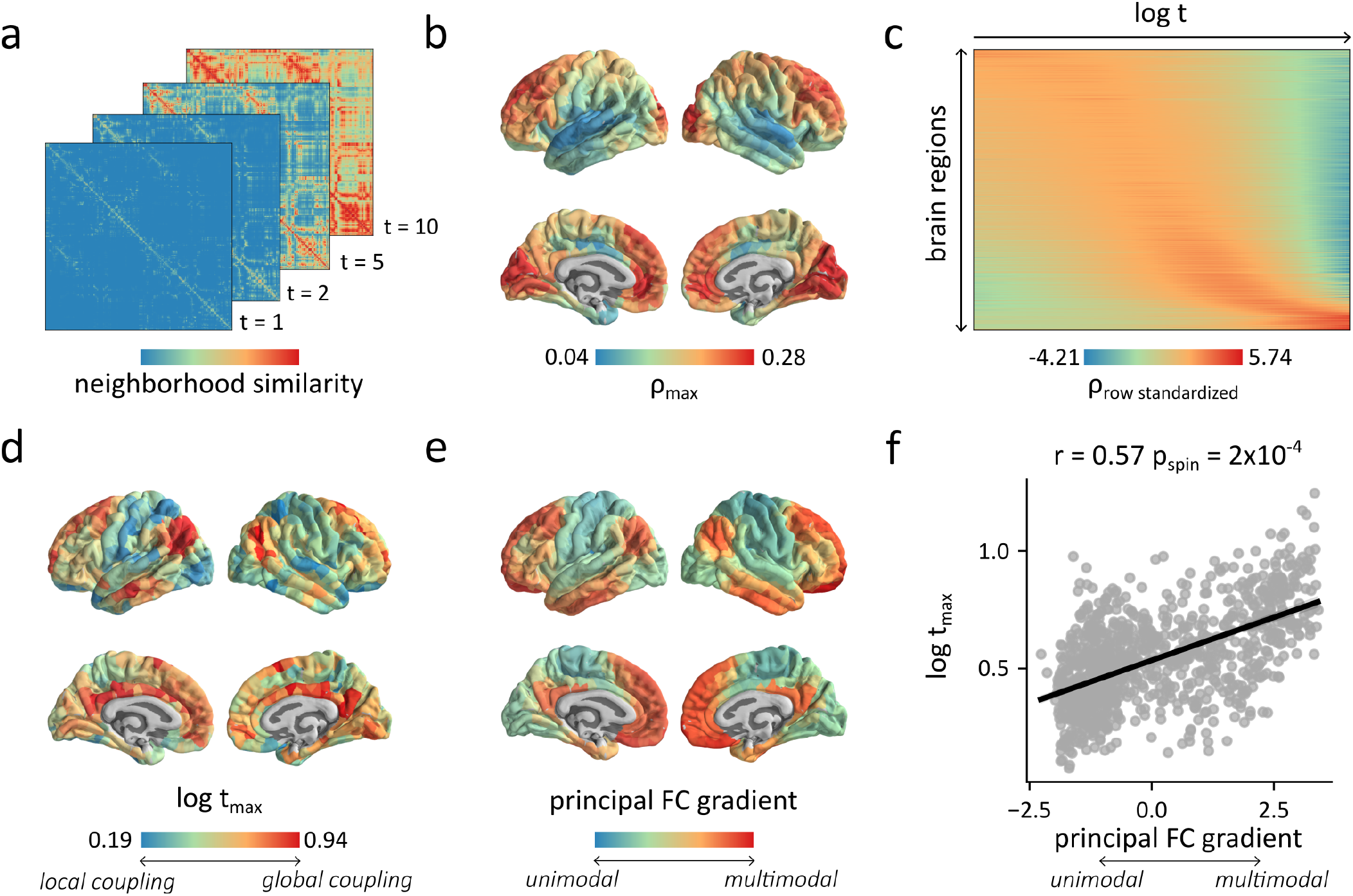
Multiscale structure-function coupling. **(a)** Neighborhood similarity for *t*=1, *t*=2, *t*=5 and *t*=10. The neighborhood similarity between pairs of brain regions was computed as the cosine similarity between their transition probability vectors, and was computed for a range of topological scales ranging from 10^*–*0.5^ to 10^1.5^. **(b)** Maximal correlations between the neighborhood similarity profiles and functional connectivity profiles of individual brain regions, averaged across subjects. **(c)** Row-standardized heatmap of the correlations between the functional connectivity and neighborhood similarity profiles of individual brain regions, as *t* increases logarithmically. **(d)** Values of *t* at which the correlation between neighborhood similarity and functional connectivity profiles is maximal, for individual brain regions, averaged across subjects. **(e)** Topographic distribution of the first (principal) gradient of functional connectivity estimated using diffusion map embedding, which reflects the main organizational axis of the brain, ranging from primary sensory and motor regions to transmodal regions [43, 50]. **(f)** Relationship between the principal gradient of functional connectivity and log *t*_max_ (r=0.57; *p*_spin_ = 2*x*10^*–*4^).

For every subject, we then measured the Pearson correlation between the functional connectivity and the neighborhood similarity of edges with positive functional connectivity weights. Measuring this correlation for every value of *t* (Fig. S3a), we find the largest correlation at *t* = 2.69 (mean = 0.22, SD = 0.03). Fig. S3b shows the distribution of individual correlation scores for each measure. The mean of the maximal correlations between functional connectivity and neighborhood similarity was significantly larger than the mean of the correlations obtained by correlating the functional connectivity matrix to the weights of the structural network’s adjacency matrix (mean = 0.12, SD = 0.02; *p <* 10^*–*40^), the weights of the structural network’s shortest paths matrix (mean = 0.16, SD = 0.03; *p <* 10^*–*20^), and the weights of the structural network’s communicability matrix (mean = 0.16, SD = 0.02; *p <* 10^*–*25^). The mean of these correlations was also significantly larger than the mean correlation between functional connectivity and Euclidean distance (mean = 0.20, SD = 0.03; *p* = 0.00046). These results are in accordance with previous results demonstrating that incorporating multiscale processes into structure-function coupling predictions is advantageous ([1, 9]).

Importantly, the present framework does not assume that structure-function relationships are uniform across the brain, and instead opens the possibility that functional interactions occur at different scales for different brain regions. To investigate this possibility, we computed the Spearman correlations between the neighborhood similarity profiles and positive-valued functional connectivity profiles of individual brain regions. We averaged the correlations obtained across individual connectomes for each value of *t*, and identified the maximal correlation (*ρ*_max_) of every brain region (Fig. 4b). We find that the optimal scale for which this regional correlation is maximal varies considerably across brain regions (Fig. 4c).

As discussed in the previous section, the optimal communication scale of brain regions varies along a unimodal-multimodal axis. We therefore hypothesize that the scale that best captures structure-function coupling for individual regions – the scale at which the correlation between neighborhood similarity and functional connectivity is maximal – similarly varies along a unimodal-multimodal axis. Fig. 4d shows the topographic distribution of the *t* values at which the correlation between neighborhood similarity and functional connectivity is maximal (*t*_max_). We see that the pattern indeed outlines the putative unimodal-transmodal hierarchy, such that functional connectivity profiles of unimodal regions are better captured by smaller neighbourhoods (small *t*), while functional connectivity in trans-modal regions is better captured inside more extensive neighbourhoods. This relationship with the unimodal-transmodal hierarchy is highlighted by a significant correlation (*r* = 0.57, *p*_spin_ = 2 × 10^*–*4^) between the optimal values of *t* and the first (principal) gradient of functional connectivity (Figs. 4e, f). This continuous gradient, which can be estimated using diffusion map embedding, is thought to reflect the main organizational axis of the brain, ranging from primary sensory and motor regions to transmodal regions [43, 50]. The magnitude of the regional correlations (Fig. 4b) were not significantly correlated with the optimal scale of a brain region (*r* = 0.25, *p*_spin_ = 0.09) and with the first (principal) gradient of functional connectivity (*r* = 0.24, *p*_spin_ = 0.21), suggesting that structure-function coupling is not weaker in multimodal regions per se, but rather, that structure-function coupling is scale-specific.

We next compare the maximal correlations obtained with neighborhood similarity (*ρ*_ns_) to the correlations obtained with two topological measures, namely shortest path (*ρ*_sp_) and communicability (*ρ*_com_), and with a geometric measure, namely Euclidean distance (*ρ*_ed_). These measures have been previously used to study variations in local structure-function coupling [85]. Fig. S4a shows the relationship between the regional correlations obtained with neighborhood similarity and the correlations obtained by comparing functional connectivity to the other measures. The optimal correlations obtained with neighborhood similarity were generally larger than those obtained with the three measures. We next compute the difference between the correlation scores obtained with neighborhood similarity and the correlation scores obtained with the other three measures. We find that the largest local differences are observed in multimodal brain regions (Fig. S4b). Namely, for all three measures, neighborhood similarity was relatively better at predicting functional connectivity in multimodal brain regions. For all three measures, we found a significant correlation between the first (principal) gradient of functional connectivity and the local differences (Fig. S4c). Altogether, these results demonstrate that while other measures perform as well (or better) in the prediction of functional connectivity in sensory regions, neighborhood similarity is significantly better at predicting functional connectivity in multimodal brain regions.

### Sensitivity and replication

We ultimately asked if the results are sensitive to different processing choices, if they are replicable with different parcellations and if they are replicable in an independently acquired dataset. In the present report, we delineated local topological neighborhoods using unbiased random walks. To ensure that the observed results are not dependent on our choice of this particular dynamical process, we repeated the analyses using personalized PageRank vectors (i.e. random-walks with restarts) and the normalized Laplacian matrix (i.e. diffusion process). These alternative dynamical processes also allow for multiscale investigations with a parameter that can be tuned to constrain their length (see *Methods* for more details). We also repeated the analyses using binarized networks, and to ensure that the log transformation of the streamline densities did not bias the results, we replicated all experiments using the streamline densities scaled to values between 0 and 1 as the weights of the structural connections (instead of their log-transform). To ensure that the results do not depend on parcellation resolution [92], we replicated all experiments with the same dataset, but parcellated into 114, 219 or 448 cortical brain regions. Finally, to ensure that the results were replicable in an independently acquired dataset, we repeated our analyses in a *Validation* dataset (HCP, N=201), which was parcellated according to a functional parcellation of 800 nodes [69]. We obtain similar results for all sensitivity and replication experiments. The optimal communication scales are presented in Fig. S5 and Fig. S6 shows their relationship with the *t*_opti_ values presented in the main text. Fig. S7 shows the topographic distributions of *t*_max_ values, and Fig. S8 shows that those distributions are significantly correlated with the principal FC gradient. Finally, Fig. S9 shows that as the parcellations get more fine-grained, structure-function coupling must be predicted by considering dynamical processes unfolding in larger neighborhoods.

## DISCUSSION

In the present report, we study how inter-regional communication between brain regions occurs over multiple topological scales. By tracing the trajectory of a region’s closeness in expanding neighborhoods, we identify topological attributes that mediate transitions from more localized communication to more global communication. We find that less diverse unimodal regions show preference for local communication and more diverse multimodal regions show preferences for global communication. These preferences manifest as scale-specific structure-function relationships with the functional connectivity of unimodal regions emerging from local communication in small-scale circuits and the functional connectivity of multimodal regions emerging from global communication in large-scale, poly-synaptic, circuits.

Numerous reports have found evidence of regional differences in centrality measures [16, 30, 38, 74, 82, 94, 96]. These studies were however performed at a single scale, eschewing the possibility that communication occurs across a spectrum of local, intermediate and global scales [4]. In other words, traditional methods overlook the possibility that proximal populations with similar functions engage in a different mode of communication from more distant populations with dissimilar functions. By tracing the trajectory of a node’s centrality over a spectrum of local neighborhoods, we show how brain regions communicate over multiple topological scale, and therefore naturally reveal their functional diversity.

The diversity of a brain region is typically estimated from the number of direct connections within-vs. between-modules [35, 64]. Regions with diverse connection profiles, with links to many specialized communities, are theoretically well-placed to integrate information from multiple domains [6–8, 66, 94]. By considering interactions over multiple hops, we not only characterize the diversity of a node’s direct connections, but also the diversity of its higher-order relationships with other regions. Moreover, our method allows for the characterization of functional diversity across a continuous range of scales, eliminating the need to partition the network into communities. This property may prove to be methodologically convenient and theoretically desirable for two reasons. First, brain networks possess prominent community structure at multiple scales, and there may not exist a single “characteristic” scale [10, 11]. Second, the community structure of the brain may not be exclusively assortative [13, 26, 63], yet most community detection algorithms assume the presence of assortative communities [28]. By tracking functional diversity over a range of expanding neighbourhoods, we avoid having to make assumptions about the presence, nature or scale of communities.

By identifying the optimal scale at which a brain region can communicate with neighboring regions, we find that regions participating in fewer integrative functions, such as primary visual, auditory and somatosensory cortices, optimally communicate at local scales. Conversely, polysensory regions in association or transmodal cortex optimally communicate at global scales. In other words, we show that a region’s functional specialization naturally emerges from its anatomical connectivity [52, 62]. Our results are naturally intertwined with the concept of segregation and integration in the brain [73]: local connectivity among regions with similar functions promotes specialized information processing whereas global connection patterns among regions with dissimilar functions promote integrative information processing. By tracing the trajectory of a brain region across multiple topological scales, we highlight a continuous gradient of localized versus distributed processing [44].

This localized-distributed gradient is further highlighted by local variations in structure-function coupling. The similarity of local structural neighborhoods best predicts functional connectivity in sensory areas, suggesting that interactions among these regions unfold mainly over local, small-scale neighbourhoods. Conversely, functional connectivity in multimodal brain regions is best predicted by the topological similarity of large-scale neighborhoods, suggesting that interactions among these regions unfold mainly over more extensive, large-scale neighbourhoods. These results build upon recent reports showing that structure-function coupling is region specific [5, 22, 67, 75, 85, 89]. Our findings suggest that this phenomenon is explained by the increasing scales at which communication processes unfold. Functional connectivity in sensory regions is easier to predict from structural connectivity because it is mediated by direct communication on the structural connectome, while functional connectivity in multimodal region is harder to predict from structural connectivity because it is mediated via more indirect, polysynaptic communication pathways [9].

Our results are in line with an emerging literature emphasizing large-scale gradients of cortical organization [31, 39, 43, 50, 60]. Our findings offer a possible explanation for how these large-scale gradients emerge from the brain’s structural embedding. Namely, at individual topological scales, areas may appear to preferentially form connections with a subset of others areas, manifesting as specific communities. As we zoom across multiple scales, however, we reveal a layered organization of interdigitated connections among areas, yielding an organizational axis of scale-specific organizational characteristics, including centrality and connection diversity. It is noteworthy that, from a geometric perspective, unimodal brain regions tend to have preferentially short-distance connections and multimodal regions have preferentially long-distance connections [57, 58, 71]. Our results are therefore consistent with recent theories suggesting that the main role of long-distance connections is to enhance the functional diversity of a brain region [12].

The present findings should be interpreted with respect to several important methodological considerations. We focused on the topological organization of networks reconstructed from diffusion-weighted imaging data using computational tractometry. This approach is prone to systematic false positives and false negatives [49, 76]. In addition, the connectomes generated are undirected, naturally limiting inferences about causal influence. We did, however, ensure that our results did not trivially depend on confounding factors by replicating our results using (1) different dynamical processes to generate neighborhoods, (2) different weights for our structural connectivity matrices, (3) a different parcellation resolution and (4) an independent validation dataset.

Altogether, the present findings demonstrate that exclusively considering communication at the global level might obscure functionally relevant features of brain networks. By studying regional embedding across multiple topological scales, we reveal a continuous range of communication preferences. In doing so, we take a step towards conceptually linking long-standing ideas in neuroscience such as integration and segregation, functional hierarchies and connection diversity.

## METHODS

### Network reconstruction

All analyses were performed in two independently collected and preprocessed datasets, one collected at the Lausanne University Hospital (*N* = 67; *Discovery*) [34] and one as part of the Human Connectome Project S900 release (*N* = 201; *Validation*) [83].

#### Discovery dataset

The *Discovery* data acquisition protocol comprised a diffusion spectrum imaging (DSI) sequence and a resting state functional magnetic resonance imaging (rs-fMRI) sequence. The *N* = 67 participants were scanned in a 3-Tesla MRI Scanner (Trio, Siemens Medical, Germany) using a 32-channel head coil. The protocol included a magnetization-prepared rapid acquisition gradient echo (MPRAGE) sequence sensitive to white/gray matter contrast (1mm in-plane resolution, 1.2mm slice thickness), a DSI sequence (128 diffusion-weighted volumes and a single b0 volume, maximum b-value 8,000 s/mm^2^, 2.2 x 2.2 x 3.0 mm voxel size), and a gradient echo-planar imaging (EPI) sequence sensitive to blood-oxygen-level-dependent (BOLD) contrast (3.3 mm in-plane resolution and slice thickness with a 0.3-mm gap, TR 1,920 ms, resulting in 280 images per participant). Initial signal processing for all MPRAGE, DSI and rs-fMRI data was performed using the Connectome Mapper pipeline [21]. Gray and white matter were segmented from the MPRAGE volume using freesurfer [23]. Grey matter was parcellated into either 114, 219, 448 or 1000 equally sized parcels [18].

Structural connectivity matrices were reconstructed for individual participants using deterministic streamline tractography on reconstructed DSI data, initiating 32 streamline propagations per diffusion direction, per white matter voxel. Within each voxel, the starting points were spatially random. For each starting point, a fiber growth continued along the ODF maximum direction that produces the least curvature for the fiber. Fibers were stopped if the change in direction was greater than 60 degrees/mm. The process was complete when both ends of the fiber left the white matter mask. The weights of the edges correspond to the log-transform of the streamline density, scaled to values between 0 and 1.

fMRI volumes were corrected for physiological variables, including regression of white matter, cerebrospinal fluid, as well as motion (three translations and three rotations, estimated by rigid body co-registration). BOLD time series were then subjected to a lowpass filter (temporal Gaussian filter with full width half maximum equal to 1.92 s). The first four time points were excluded from subsequent analysis to allow the time series to stabilize. Motion “scrubbing” was performed as described by [65]. Functional connectivity matrices were constructed by computing the zero-lag Pearson correlation coefficient between the fMRI BOLD time series of each pairs of brain regions.

#### Validation dataset

The *Validation* data acquisition protocol included a high angular resolution diffusion imaging (HARDI) sequence and four resting state fMRI sessions. All analyses were performed in a subset of *N* = 201 unrelated participants. The participants were scanned in the HCP’s custom Siemens 3T “Connectome Skyra” scanner. Further information regarding the acquisition protocol is available at [83] while more information regarding the preprocessing and the network reconstruction is available at [61].

Briefly, the dMRI data was acquired with a spin-echo EPI sequence (TR=5,520 ms; TE=89.5 ms; FOV=210 × 180 mm ^2^; voxel size=1.25 mm ^3^; b-value=three different shells i.e., 1,000, 2,000, and 3,000 s/mm ^2^; number of diffusion directions=270; and number of b0 images=18) and the rs-fMRI data was acquired using a gradient-echo EPI sequence (TR=720 ms; TE=33.1 ms; FOV=208 × 180 mm ^2^; voxel size=2 mm ^3^; number of slices=72; and number of volumes=1,200). The data was pre-processed according to the HCP minimal preprocessing pipelines [29].

Structural connectomes were reconstructed from the dMRI data using the MRtrix3 package [77]. Fiber orientation distributions were generated using the multishell multi-tissue constrained spherical deconvolution algorithm from MRtrix [24, 46]. The initial tractogram was generated with 40 million streamlines, with a maximum tract length of 250 and a fractional anisotropy cutoff of 0.06. Spherical-deconvolution informed filtering of tractograms (SIFT2) was applied to reconstruct whole brain streamlines weighted by cross-section multipliers [72]. To ensure that our results were not confounded by the parcellation scheme or resolution, grey matter was parcellated with a different parcellation. This time, grey matter was parcellated into 800 cortical regions according to the Schaefer functional atlas [69]. Functional connectivity matrices were constructed for individual subjects by computing the zero-lag Pearson correlation coefficient between the fMRI BOLD time series of each pairs of brain regions. The functional weights of the four resting-state sessions were then averaged for each individuals.

### Yeo intrinsic networks and von Economo classes

To facilitate our analyses, nodes of the brain networks were stratified according to their membership to seven intrinsic functional networks and seven cytoarchitectonic classes. The seven intrinsic functional networks were identified by applying a clustering technique on restingstate fMRI data from 1000 subjects. More details can be found in [91]. The seven resting-states parcellation, in the FreeSurfer fsaverage5 surface space, was first downloaded from https://github.com/ThomasYeoLab/CBIG/. We then attributed to each parcel of the 1000 nodes Cammoun parcellation the most common intrinsic network assignments of its vertices. The seven cytoarchitectonic classes consist in an extended version of the classical von Economo atlas [87, 88]. The class of each parcel was manually assigned based on visual comparison with the von Economo and Koskinas’s parcellation and anatomical landmarks [84, 86].

### Multiscale topological neighborhoods

Local neighborhoods were characterized by modelling dynamical process initiated from individual nodes in the networks. By controlling the length of these processes, we controlled the topological size of the delineated neighborhoods. Our main results relied on unbiased random-walks. We further replicated our results using random walks with restarts and a heat-diffusion process.

#### Unbiased random-walks

Given an adjacency matrix **A** where *A*_*ij*_ corresponds to the weight of the edge connecting nodes *i* and *j*, the probability that a walker at node *i* transitions to node *j* in a single iteration is given by 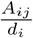, where *d*_*i*_ is the degree (strength if weighted) of node *i* and corresponds to the sum of the weights of the edges leaving node *i*. The overall transition probabilities of a network can be represented in a transition matrix **P** such that:

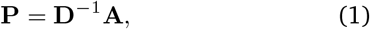

where **D** is the diagonal matrix with the value *D*_*ii*_ corresponding to the degree of node *i*. Given an initial distribution of random walkers **p**(0), it is possible to compute the distribution of these random walkers at a discrete time *t*, **p**(*t*):

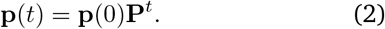

The vector **p**(*t*) indicates the proportion of walkers located at any other node at time *t*. We initiate this random walk process on a single node *i* by setting the initial distribution **p**(0) to be equal to 0 everywhere, and be equal to 1 in position *i*. This initial vector can be written as **e**_*i*_. The transition probability vector at time *t*, given that the random walk process was started on node *i* can then we written as:

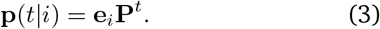

The discrete-time random walk process defined above can be “continuized” by considering the interval of time between two moves as an exponential random variable with *λ* = 1 [54]. The transition probability vector, for a random walk process initiated on node *i*, is then given by:

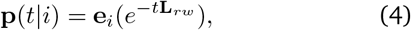

where **L**_*rw*_ = **I – P** is the graph random-walk normalized Laplacian.

By initiating a random-walk process from a single node, we can measure the topological proximity between this node and the other nodes in the network. By increasing the value of *t*, we increase the length of the random-walks, and consequently measure the topological proximity of nodes in larger neighborhoods. We measured the transition probabilities for 100 time scales *t*, logarithmically spaced between 10^*–*0.5^ and 10^1.5^.

#### Random-walks with restarts

A variant of this equation consists in adding to this random-walk process an additional probability that the random walker randomly teleports itself back to the seed node. Random walks with teleportation are the basis for the PageRank algorithm [15] and ensure that random walks on directed networks do not get trapped in absorbing states. The equation, given that the random walk process is initiated on a single node *i*, is defined as follows:

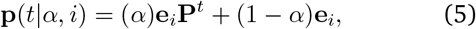

where *α* corresponds to the damping factor. By varying the damping factor, we can decrease the probability of restart of the random walks and therefore increase their length. More specifically, the probability that a random-walks is of length *l* (*Prob*[*L* = *l*]) is geometrically related to the value of the parameter *α*:

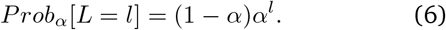

Ultimately, the stationary distribution of this Markov chain, which can be computed using the power iteration method, indicate the probability that random walkers of a certain length, starting from a single source node, reach other nodes in the network. We computed the stationary distribution, also known as the personalized PageRank, for 99 values of *α*, linearly spaced between 0.01 and 0.99.

#### Diffusion

An alternative method to dynamically measure the topological proximity of brain regions in a network consists in modelling a heat-diffusion process on the network. More specifically, the Laplacian matrix (**L**) of the network is used to compute the distribution of some material at time *t*, given that the process was initiated at node *i*:

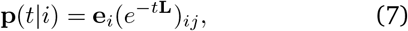

where

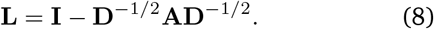

Again, by varying the value of *t*, we can vary the size of the neighborhoods on which the diffusion process unfolds, and therefore measure the topological proximity of nodes in the network given dynamical processes unfolding at increasingly large scales. We measured the transition probabilities for 100 time scales *t*, logarithmically spaced between 10^*–*0.5^ and 10^1.5^.

### Multiscale closeness centrality

Measures of network centrality often consider a node’s relationship with all of the other nodes. For instance, the closeness centrality of a node in a network can be measured by computing the inverse of its averaged topological distance to the other nodes in the network:

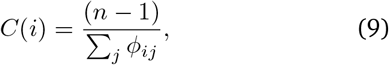

where n corresponds to the number of nodes in the network parcellation and *ϕ*_*ij*_ corresponds to the weighted shortest path between nodes *i* and *j*.

To capture the centrality of a node at different scales, we propose a new measure. This measure consists in computing the multiscale closeness centrality of a node as a weighted average using the probability vector **p** as a weight function prioritizing the node’s relationships with nodes that are topologically close over nodes that are topologically remote. Specifically, given a scale-dependent weight vector **p**(*t*|*i*), the multiscale closeness centrality of a node *i* for the specified scale *t* is defined as:

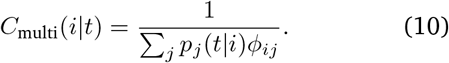

The topological distance between a pair of connected nodes *i* and *j* was measured as the inverse of the connection weight between the two nodes, and the topological shortest paths between pairs of nodes were subsequently retrieved using the Dijkstra’s algorithm. The code to compute this multiscale measure of closeness centrality is available in the project’s github repository (https://github.com/netneurolab/bazinet_multiscale)

### Clustering coefficient

The clustering coefficient of a node corresponds to the number of triangles attached to it, normalized by the maximum number of possible triangles that could be attached to the node [90]:

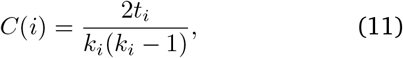

where *k*_*i*_ is the degree of node *i* and *t*_*i*_ is the number of triangles attached to node *i*. A weighted version of the clustering coefficient, which can be viewed as a measure of the average “intensity” of triangles around a node, can also be expressed as follows [59]:

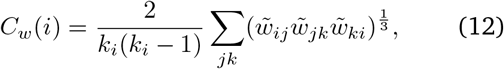

where 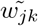 is the weight of the connection between nodes *j* and *k*, divided by the largest weight in the network. The clustering coefficients were computed using the Brain Connectivity Toolbox (https://github.com/aestrivex/bctpy) [68].

### Community detection

Communities are groups of nodes with dense connectivity among each other. The Louvain method was used to identify a community assignment or partition that maximizes the quality function *Q* [14]:

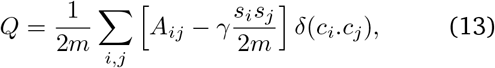

where *A*_*ij*_ is the weight of connection between nodes *i* and *j, s*_*i*_ and *s*_*j*_ are the directed strengths of *i* and *j, m* is a normalizing constant, *c*_*i*_ is the community assignment of node *i* and the Kronecker *δ*-function *δ*(*u, v*) is defined as 1 if *u* = *v* and 0 otherwise. The resolution parameter *γ* scales the importance of the null model and effectively controls the size of the detected communities: larger communities are more likely to be detected when *γ <* 1 and smaller communities (with fewer nodes in each community) are more likely to be detected when *γ >* 1.

To detect stable community assignments for our structural connectomes, we first constructed a consensus network from the individual connectomes. This network was generated such that the mean density and edge length distribution observed across individual participants was preserved [12, 55, 56]. The edges were weighted as the average weight across individual networks for which these edges existed. Using this consensus network, we initiated the algorithm 100 times at each value of the resolution parameter and consensus clustering was used to identify the most representative partitions [48]. This procedure was repeated for a range of 100 resolutions between *γ* = 0.25 and *γ* = 7.5. We then quantified the similarity between pairs of consensus partitions using the *z* score of the Rand index [78]. We next identified five values of *γ* at which the generated partitions showed high mutual similarity and persisted through stretches of *γ* values. This procedure yielded partitions of 4, 6, 9, 12 and 16 communities (corresponding to *γ*=0.54, 1.08, 2.06, 2.85, 7.16). The whole procedure was implemented using code available in the netneurotools python toolbox (https://github.com/netneurolab/netneurotools).

### Participation coefficient

Given a partition, we quantify the diversity of a node’s connections to multiple communities using the *participation coefficient* [35]. The participation coefficient is defined as

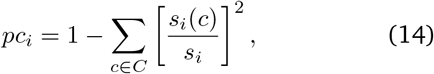

where *s*_*i*_ is the total strength of node *i, s*_*i*_(*c*) is the strength of *i* in community *c* and the sum is over the set of all communities *C*. Nodes with a low participation coefficient are mainly connected with nodes in a single community, while nodes with a high participation coefficient have a diverse connection profile, with connections to multiple communities. The participation coefficients were computed using the Brain Connectivity Tool-box (https://github.com/aestrivex/bctpy) [68].

### Autocorrelation-preserving permutations

To assess the significance between the principal functional connectivity gradient and *t*_max_, we relied on autocorrelation-preserving permutations and generated null distributions that preserve the spatial autocorrelation of the original brain map. By preserving this autocorrelation, we ensure that the null distributions do not violate the assumption of exchangeability and that permutation tests will not generate inflated p-values [2, 51]. To generate autocorrelation-preserving permutations, we first created a surface-based representation of the Cammoun atlas on the FreeSurfer fsaverage surface using the Connectome Mapper toolkit (https://github.com/LTS5/cmp, [21]). We identified the vertices closest to the center-of-mass of each parcel and used the spherical projection of the fsaverage surface to define spherical coordinates for each parcel. We then applied randomly-sampled rotations to the spherical atlas and reassigned each parcel to the closest parcel following this rotation. Each rotation was applied to one hemisphere and then mirrored to the other hemisphere. This process was repeated 10000 times using code available in the netneurotools python toolbox (https://github.com/netneurolab/netneurotools). The empirical distributions were then compared to these spatially-autocorrelated nulls and two-sided p-values (*p*_spin_) were computed.

### Neighborhood similarity

The pairwise neighborhood similarity of nodes in the network, for a particular scale *t* can be represented in a matrix **S**(*t*), where *S*_*ij*_(*t*) corresponds to the cosine similarity between the transition probability vectors of nodes *i* and *j*, for the scale *t*:

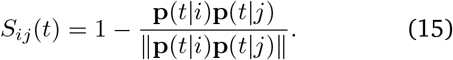

The code to compute neighborhood similarity is available in the project’s github repository (https://github.com/netneurolab/bazinet_multiscale).

### Communicability

For a binary adjacency matrix **A**, communicability is defined as

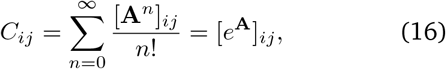

with walks of length *n* normalized by *n*!, ensuring that shorter, more direct walks contribute more than longer walks [25]. For a weighted adjacency matrix, this definition can be extended as

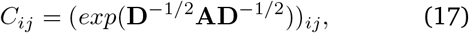

where **D** is the diagonal degree matrix [20]

### Principal FC gradient

The principal gradient of functional connectivity is thought to reflect the main organizational axis of the brain, ranging from primary sensory and motor regions to transmodal regions [43, 50]. This gradient can be reconstructed using diffusion map embeddding, a nonlinear dimensionality reduction algorithm [19]. The algorithm seeks to project a set of embeddings into a lower-dimensional Euclidean space. Briefly, the similarity matrix among a set of points (in our case, the correlation matrix representing functional connectivity) is treated as a graph, and the goal of the procedure is to identify points that are proximal to one another on the graph. In other words, two points are close together if there are many relatively short paths connecting them. A diffusion operator, representing an ergodic Markov chain on the network, is formed by taking the normalized graph Laplacian of the matrix. The new coordinate space is described by the eigenvectors of the diffusion operator.

We set the diffusion rate *α* = 0.5. The eigenvalues *λ* were divided by 1 – *λ* to provide noise robustness and eliminate the need for a diffusion time (t) parameter. For each dataset and parcellation, the principal gradient was computed on a consensus functional connectivity matrix computed by averaging the pairwise correlations obtained across individuals and setting the negative values to 0. The procedure was implemented using the mapalign Toolbox (https://github.com/satra/mapalign).

### Generative model

A stochastic block model [42] was used to generate an artificial hierarchically modular network of 2000 nodes with a layer of 4 equally-sized communities and another of 2 equally-sized communities. The edge probability matrix *P* was defined as follows:

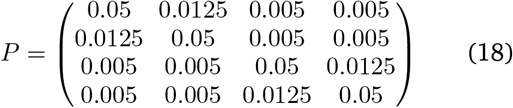

The stochastic block model was implemented using the NetworkX package [37]. The two-dimensional embedding of this network was generated using the ForceAtlas2 algorithm [45].

## Data availability

The *Discovery* dataset (Lausanne) is available at https://doi.org/10.5281/zenodo.2872624 and the *Validation* dataset (Human Connectome project) is available at https://www.humanconnectome.org/study/hcp-young-adult. The code used to conduct the analyses presented in this paper is available at https://github.com/netneurolab/bazinet_multiscale.

## ACKNOWLEDGMENTS and DISCLOSURES

We thank Laura Suarez, Justine Hansen, Golia Shafiei, Bertha Vazquez-Rodriguez, Ross Markello and Zhen-Qi Liu for insightful comments. VB acknowledges support from the Fonds du Recherche Québec - Nature et Technologies. BM acknowledges support from the Natural Sciences and Engineering Research Council of Canada (NSERC Discovery Grant RGPIN #017-04265), from the Canada Research Chairs Program and from the Canada First Research Excellence Fund, awarded to McGill University for the Healthy Brains for Healthy Lives initiative.

---

**Figure S1.**
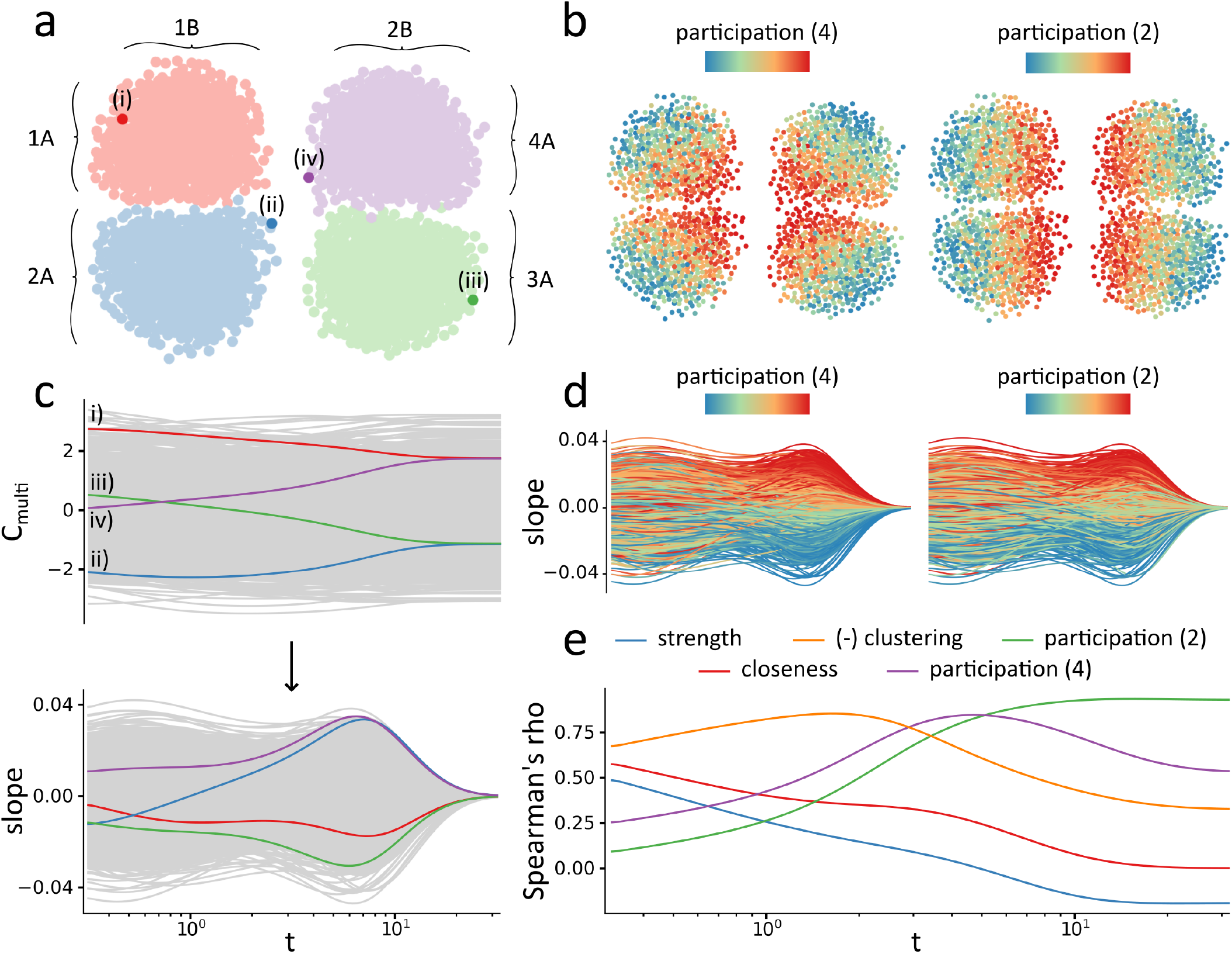
Closeness trajectories in a modular network. In this figure, we illustrate how the closeness trajectory of a node in a network reflects its functional diversity. We first generated an artificial network of n=2000 nodes using a stochastic block model **(a)** Two-dimensional representation, generated using the ForceAtlas2 algorithm [45], of the hierarchical modular network. Its community structure is composed of two large communities of 1000 nodes (1B and 2B) as well as four smaller communities of 500 nodes (1A, 2A, 3A and 4A). **(b)** The functional diversity of individual nodes is usually characterized by the participation coefficient, which compares the number of within- and between-community connections [35]. By measuring the participation coefficient for a 4-communities partition (left) and for a 2-communities partition (right), we can identify nodes in the network that are important for communication between the four sub-communities, and between the two larger communities. Nodes with large participation coefficients are located at the borders between the communities. **(c)** Two nodes can have the same global centrality, yet different local architectures. For instance node (i) in the red community and node (iv) in the purple community have the same global closeness centrality, but their location in the low dimensional embedding illustrated in panel (a) suggests that they have very different connection profiles. Node (i) is located on the external edge of the red community while node iv) is located on the internal edge of the purple community, closer to the border between the four communities. Their different connection diversity is captured by their different closeness centrality trajectories. Node (iv) shows an increase in its centrality as the topological scale of the probed neighborhood increases while node (i) shows a decrease in its centrality. Similar comparisons can be done between nodes (ii) and (iii). Variations in a node’s closeness can be quantified by measuring the local variations (*slope*) in their centrality. **(d)** Closeness trajectories of the network’s nodes, colored according to their participation coefficients given a 4-communities parcellation (left) and a 2-communities parcellation (right). Nodes with a large participation coefficient show a positive slope and nodes with a small participation coefficient show a negative slope. **(e)** Slopes can be measured for any value of *t*, highlighting the diversity of a node’s connections at the chosen scale. At small values of *t*, slopes are correlated with local measures of diversity, such as clustering coefficient (negative correlation), and as *t* increases, slopes are correlated with increasingly more global measures of diversity (e.g. participation coefficients).

**Figure S2.**
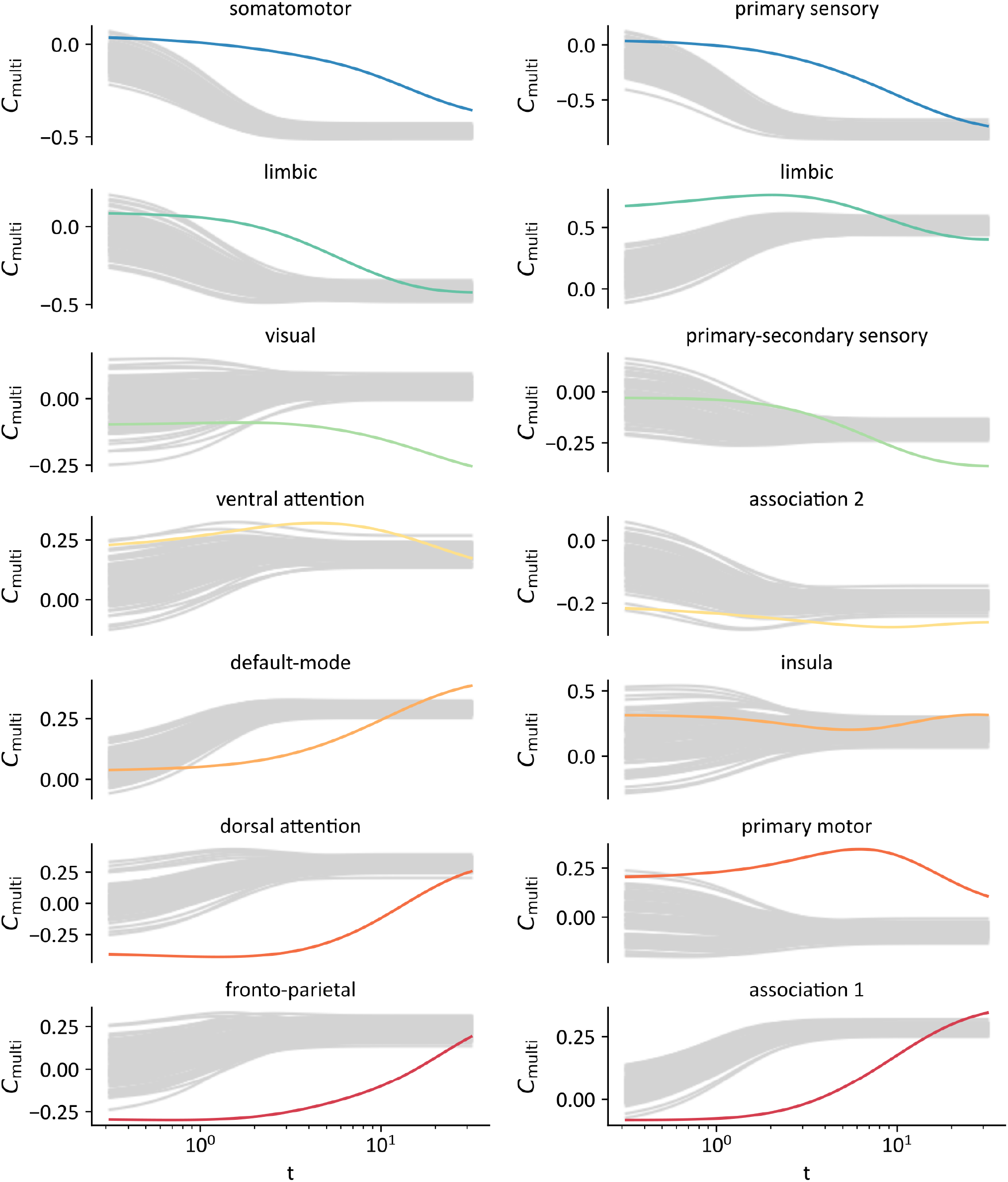
*C*_multi_ in degree-preserved randomized networks. Mean *C*_multi_ of seven intrinsic functional networks (right) and seven cytoarchitectonic classes (left) as t increases (colored lines), compared to the mean *C*_multi_ of the same intrinsic networks and cytoarchiteconic classes in 100 degree-preserving randomized networks (gray lines). The empirical *C*_multi_ scores are averaged across the individual connectomes while each individual gray line represents the *C*_multi_ of a randomized network. The surrogate networks were constructed by first generating a consensus structural connectivity network that preserved the mean density and edge length distributions of the individual networks [12, 55, 56]. The edges of the consensus network were then randomly swapped to create randomized networks with the same degree sequence and density as the original network [53].

**Figure S3.**
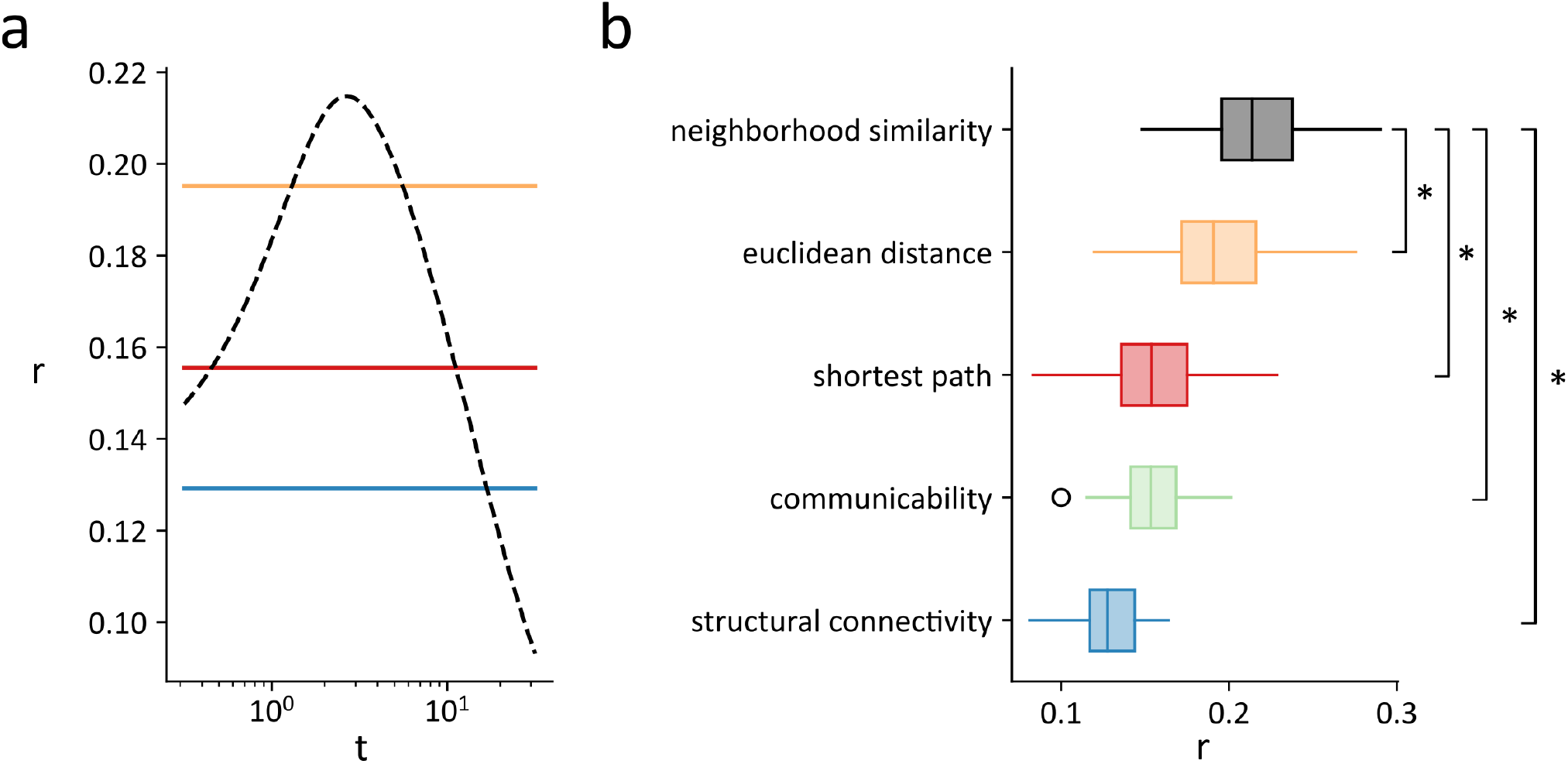
Connectome-wide structure-function coupling. **(a)** Mean, across subjects, of the Pearson correlations between functional connectivity and edge-wise measures of communication (neighborhood similarity with respect to *t*, black/dashed; structural connectivity weight, blue; communicability, green, shortest path, red; Euclidean distance, yellow). The correlation between neighborhood similarity and functional connectivity is maximal at *t* = 2.69. (**b**) Distributions of subject-wise correlations between edge-wise measures of communication and functional connectivity. The mean of the distribution of maximal correlations between neighborhood similarity and functional connectivity (mean=0.22, SD=0.03) is significantly larger than the mean of the distributions of correlations between functional connectivity and Euclidean distance (mean=0.20, SD=0.03, *p* = 4.6 × 10^*–*4^), shortest paths (mean=0.16, SD=0.03, *p <* 10^*–*20^), communicability (mean=0.16, SD=0.02, *p <* 10^*–*25^) as well as structural connectivity weights (mean=0.12, SD=0.02, *p <* 10^*–*40^).

**Figure S4.**
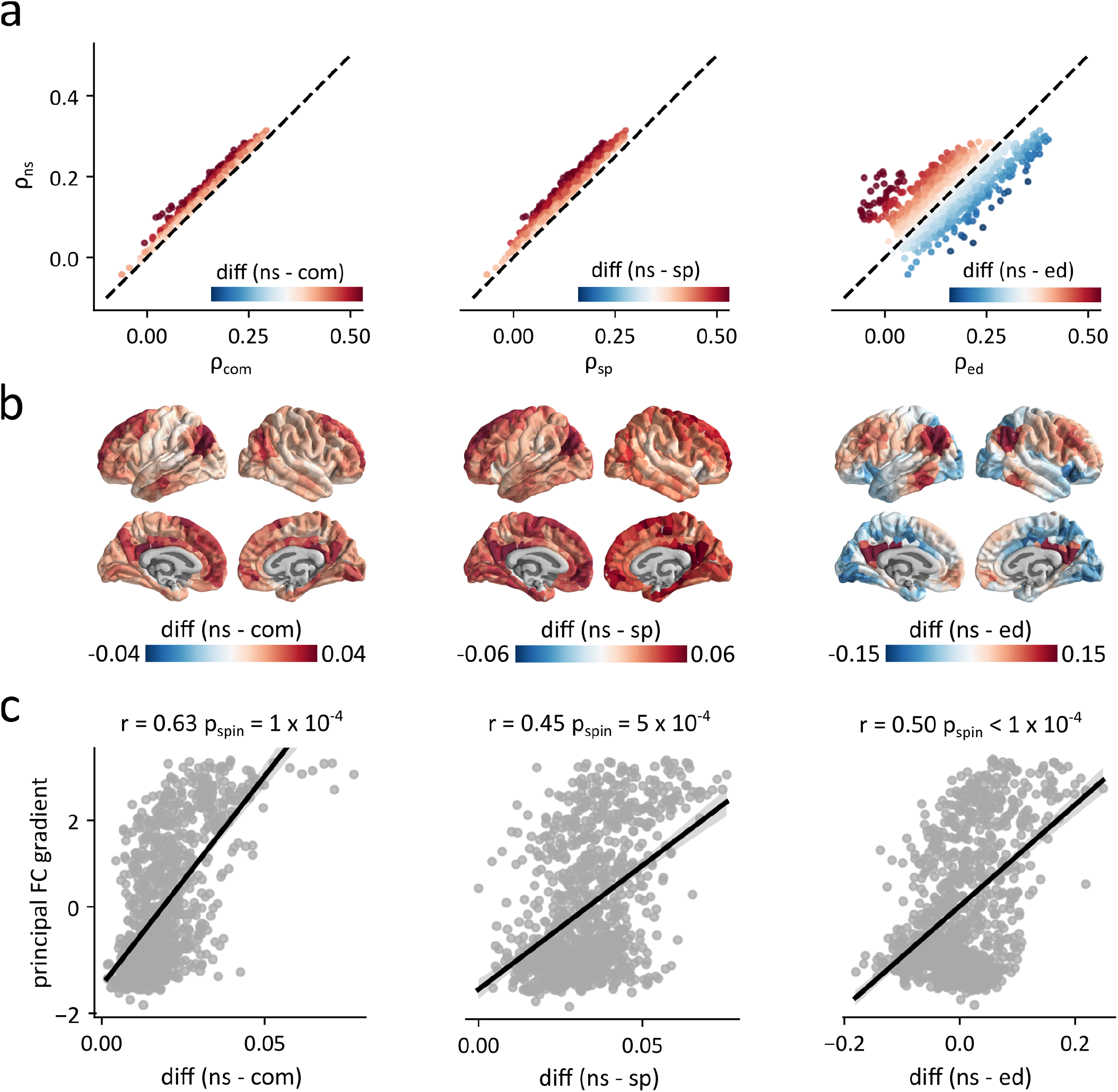
Structure-function coupling using alternate topological and geometric measures. **(a)** Relationship between the regional correlations obtained by comparing functional connectivity profiles to neighborhood similarity profiles (*ρ*_ns_) and the regional correlations obtained by comparing functional connectivity profiles to communicability (*ρ*_com_), shortest path (*ρ*_sp_) and Euclidean distance (*ρ*_ed_) profiles. The dashed black line highlights the identity line and nodes are colored according to the difference between each pair of regional correlations. (**b**) Topographic distributions of the differences between the functional connectivity correlations measured with neighborhood similarity and the functional connectivity correlations measured with communicability (left), shortest path (middle) and Euclidean distance (right). (**c**) Relationship between the correlation differences and the principal gradient of functional connectivity.

**Figure S5.**
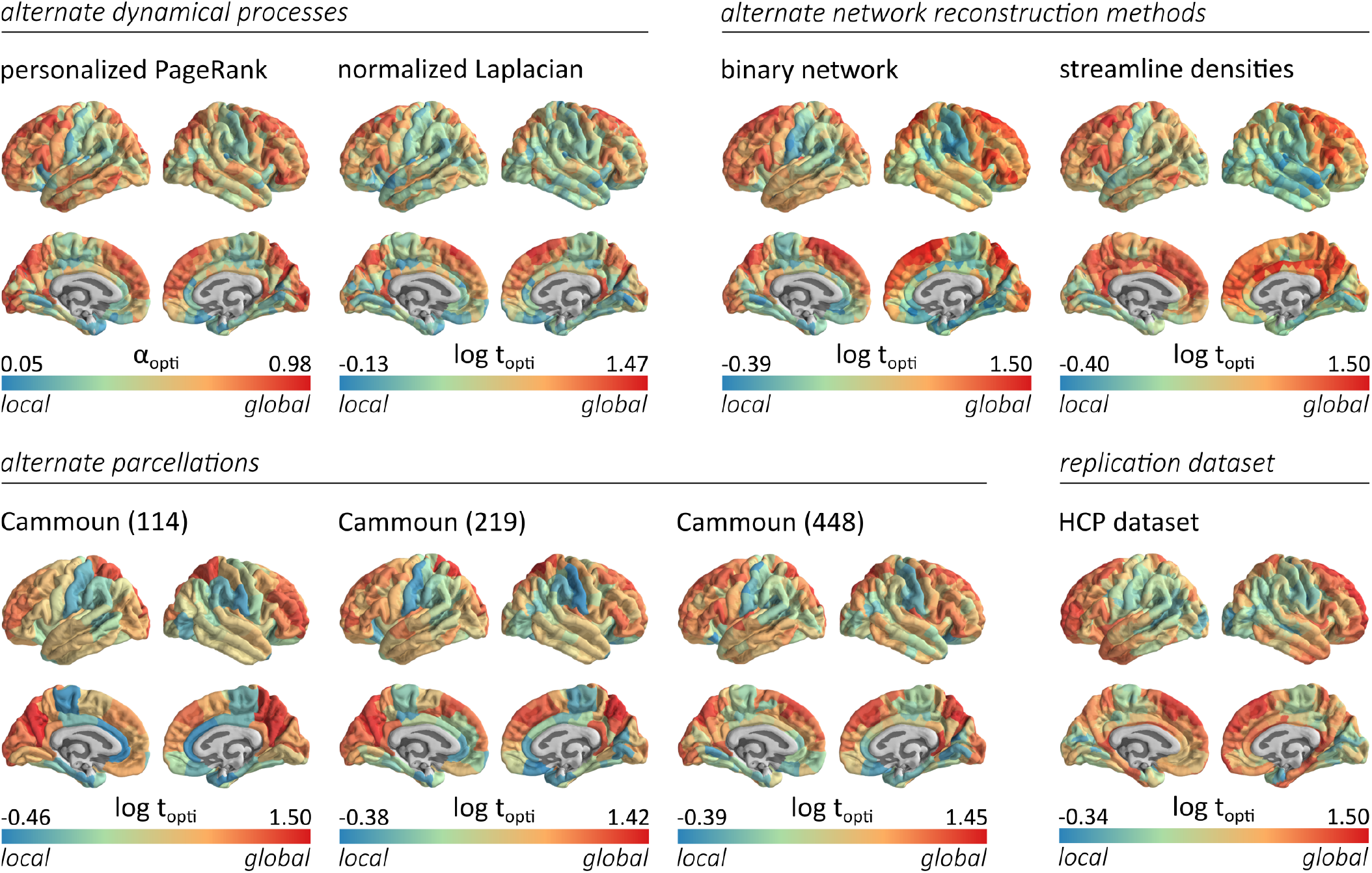
Sensitivity and replication analyses. Topographic distributions of the optimal communication scales (*α*_opti_ or log *t*_opti_), averaged across subjects, for different sensitivity or replication experiments. Top row, from left to right: Optimal closeness scales computed using personalized PageRank vectors, and using the network’s normalized graph Laplacian. Optimal closeness scales computed in the binarized structural connectomes, and in structural connectomes with weights corresponding to the streamline densities, scaled to values between 0 and 1. Bottom row, from left to right: optimal closeness scales computed in structural connectomes reconstructed using the Cammoun 114, 219 and 448 parcellations [18]. Optimal closeness scales computed in structural connectomes reconstructed from the HCP *validation* dataset. For all experiments, we find that primary sensory regions optimally communicate at local scales while association regions optimally communicate at global scales.

**Figure S6.**
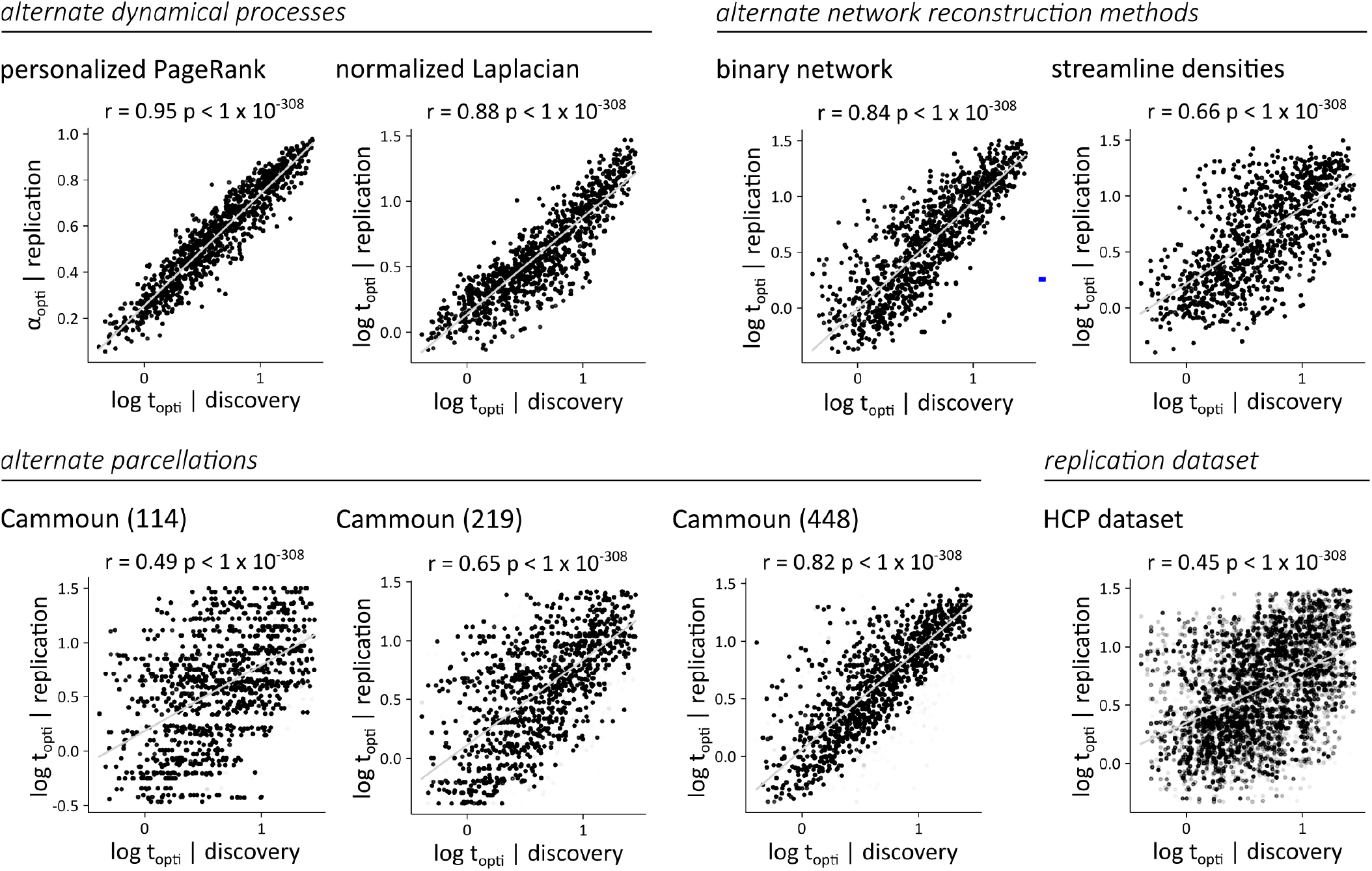
Sensitivity and replication analyses. The distributions of optimal closeness scales obtained for each sensitivity or replication test (*replication*) are compared to the distribution presented in the main text (*discovery*). The parcel-wise results are projected to the vertex level, to allow comparison between parcellations of different sizes. Top row, from left to right: optimal closeness scales computed using personalized PageRank vectors (*r* = 0.95; *p <* 1 × 10^*–*308^), and using the network’s normalized graph Laplacian (*r* = 0.88; *p <* 1 × 10^*–*308^). Optimal closeness scales computed in the binarized structural connectomes (*r* = 0.84; *p <* 1 × 10^*–*308^), and in structural connectomes with weights corresponding to the streamline densities scaled to values between 0 and 1 (*r* = 0.66; *p <* 1 × 10^*–*308^). Bottom row, from left to right: optimal closeness scales computed in structural connectomes reconstructed using the Cammoun 114 (*r* = 0.49; *p <* 1 × 10^*–*308^), 219 (*r* = 0.65; *p <* 1 × 10^*–*308^) and 448 parcellations (*r* = 0.82; *p <* 1 × 10^*–*308^) [18]. Optimal closeness scales computed in structural connectomes reconstructed from the HCP *validation* dataset (*r* = 0.45; *p <* 1 × 10^*–*308^). For all experiments (*replication*), we find significant correlations with the results presented in the main text (*discovery*).

**Figure S7.**
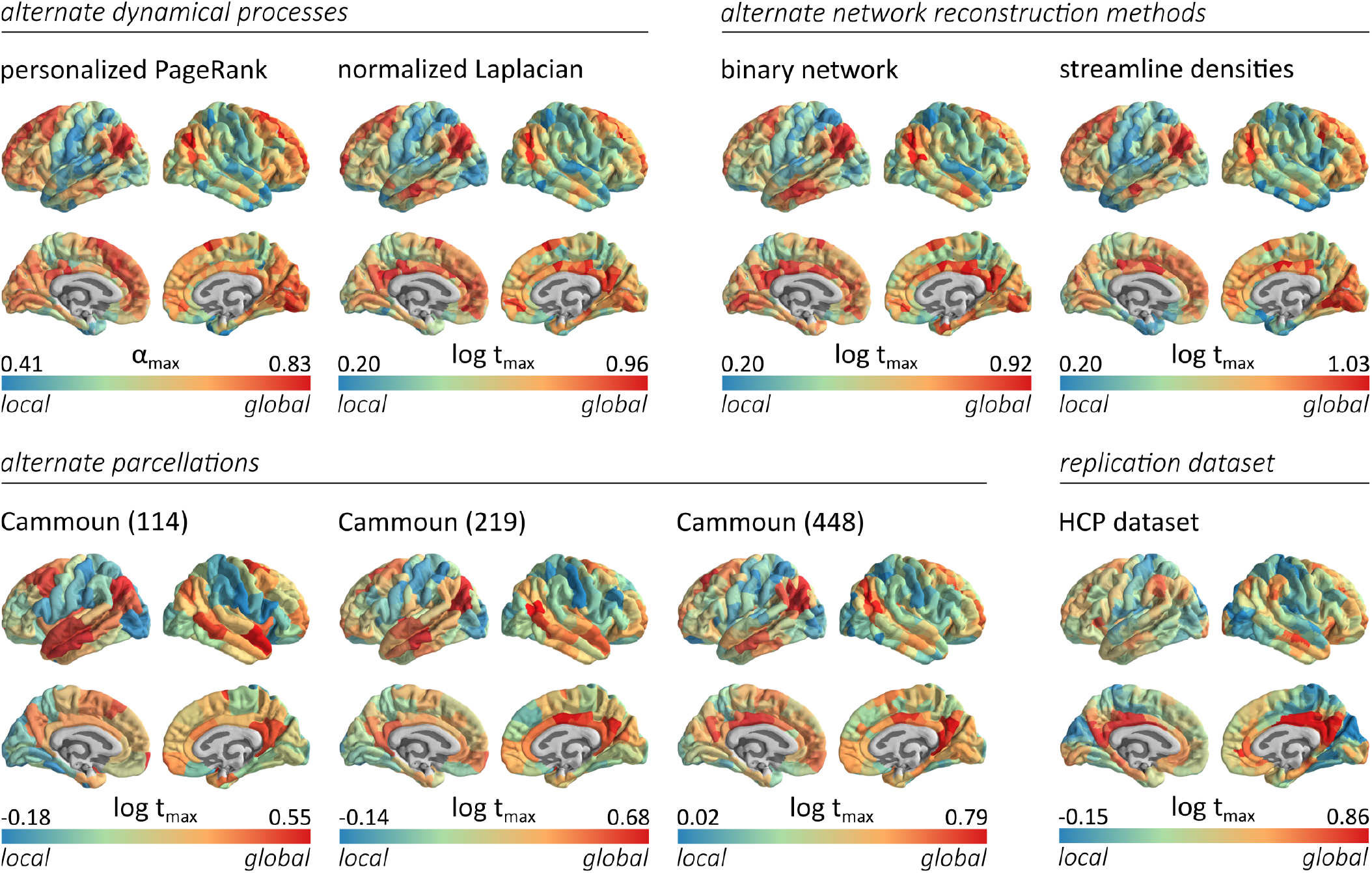
Sensitivity and replication analyses. Topographic distributions of the topological scale at which the correlation between regional functional connectivity and regional neighborhood similarity is maximal (*α*_max_ or log *t*_max_). The results are presented for different sensitivity or replication experiments. Top row, from left to right: *α*_max_ computed using personalized PageRank vectors, and log *t*_max_ computed using the network’s normalized graph Laplacian. log *t*_max_ computed in binarized structural connectomes, and log *t*_max_ computed in structural connectomes with weights corresponding to the streamline densities scaled to values between 0 and 1. Bottom row, from left to right: log *t*_max_ computed in structural connectomes reconstructed using the Cammoun 114, 219 and 448 parcellations [18]. log *t*_max_ computed in structural connectomes reconstructed from the HCP *validation* dataset. For all experiments, we observe small values of *t* in sensory regions and large values of *t* in multimodal regions. In other words, the functional connectivity of sensory regions is maximally correlated to the similarity of small, local, structural neighborhoods while the functional connectivity of multimodal regions is maximally correlated to the similarity of large-scale structural neighborhoods.

**Figure S8.**
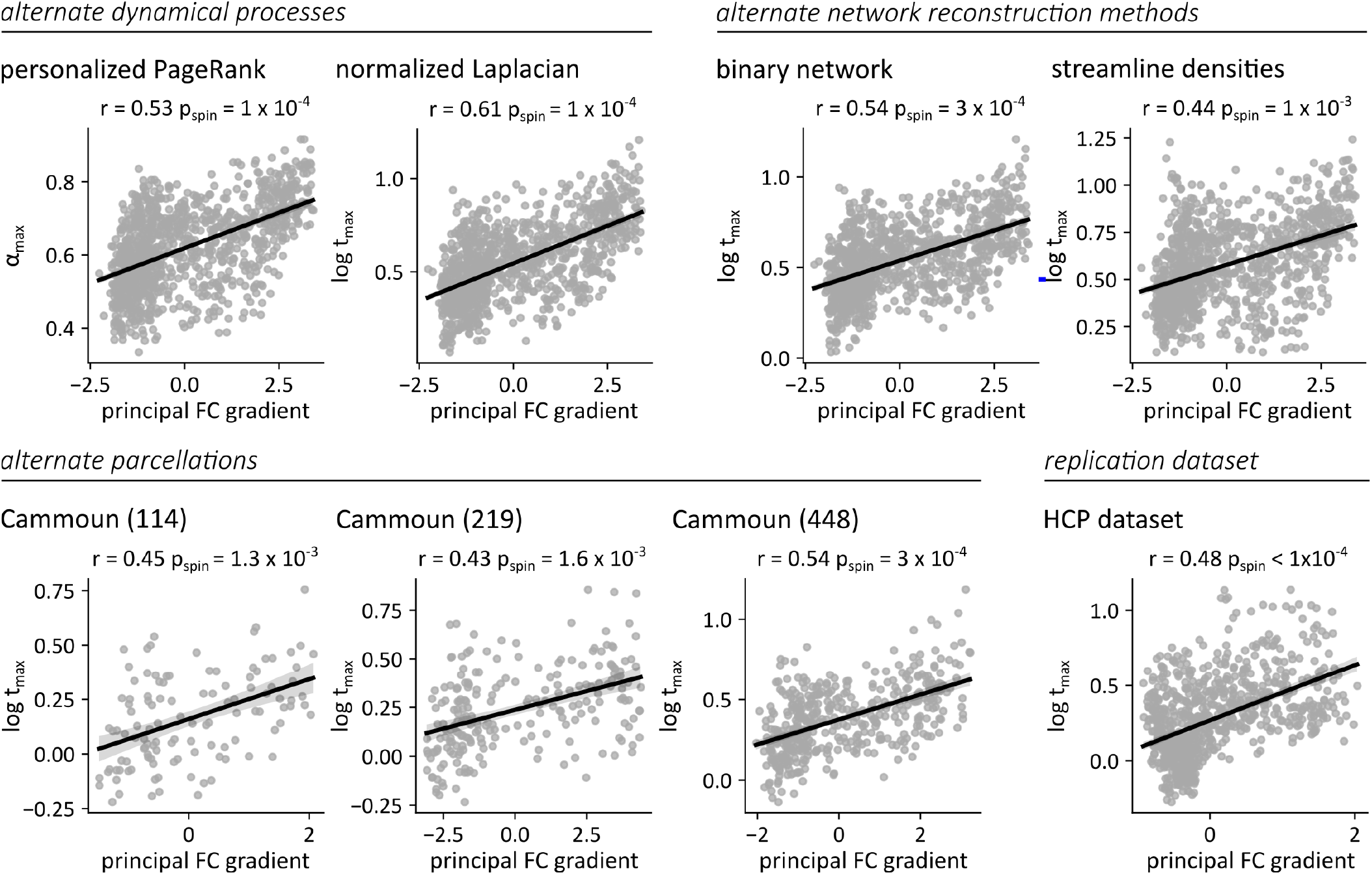
Sensitivity and replication analyses. Distributions of the topological scale at which the correlation between regional functional connectivity and regional neighborhood similarity is maximal (*α*_max_ or log *t*_max_), compared to the principal functional connectivity gradient. The results are presented for different sensitivity or replication experiments. Top row, from left to right: *α*_max_ computed using the personalized PageRank (*r* = 0.53; *p*_spin_ = 1 × 10^*–*4^), and log *t*_max_ computed using the network’s normalized graph Laplacian (*r* = 0.61; *p*_spin_ = 1 × 10^*–*4^). log *t*_max_ computed in the binarized structural connectomes (*r* = 0.54; *p*_spin_ = 3 × 10^*–*4^), and log *t*_max_ computed in the structural connectomes with weights corresponding to the streamline densities scaled to values between 0 and 1 (*r* = 0.44; *p*_spin_ = 1 × 10^*–*3^). Bottom row, from left to right: log *t*_max_ computed in structural connectomes reconstructed using the Cammoun 114 (*r* = 0.45; *p*_spin_ = 1.3 × 10^*–*3^), 219 (*r* = 0.43; *p*_spin_ = 1.6 × 10^*–*3^) and 448 (*r* = 0.54; *p*_spin_ = 3 × 10^*–*4^) parcellations [18]. log *t*_max_ computed in structural connectomes reconstructed from the HCP *validation* dataset (*r* = 0.48; *p*_spin_ *<* 1 × 10^*–*4^). For all experiments, we observe significant correlations between the principal gradient of functional connectivity and *t*_max_. In other words, these results confirm the existence of a relationship between a brain region’s coupling scale and its location in the unimodal-multimodal hierarchy.

**Figure S9.**
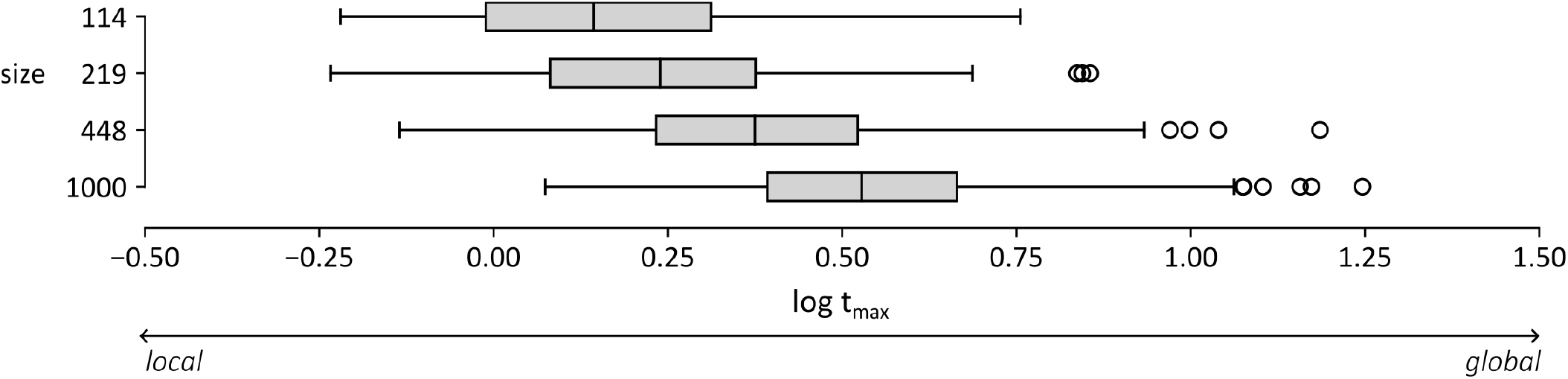
Structure-function coupling across parcellation scales. Distributions of log *t*_max_ scores, for the Cammoun 114, 219, 448 and 1000 parcellations. For fine-grained parcellations, structure-function coupling is best predicted by considering dynamical processes unfolding in larger-scale neighborhoods.

